# Identification of miRNAs and associated pathways regulated by Leukemia Inhibitory Factor in trophoblastic cell lines

**DOI:** 10.1101/410381

**Authors:** Diana M. Morales-Prieto, Emanuel Barth, Jose Martín Murrieta-Coxca, Rodolfo R. Favaro, Ruby N. Gutiérrez-Samudio, Wittaya Chaiwangyen, Stephanie Ospina-Prieto, Bernd Gruhn, Ekkehard Schleußner, Manja Marz, Udo R. Markert

## Abstract

**Introduction:** Leukemia Inhibitory Factor (LIF) regulates behavior of trophoblast cells and their interaction with immune and endothelial cells. *In vitro*, trophoblast cell response to LIF may vary depending on the cell model. Reported differences in the miRNA profile of trophoblastic cells may be responsible for these observations. Therefore, miRNA expression was investigated in four trophoblastic cell lines under LIF stimulation followed by *in silico* analysis of altered miRNAs and their associated pathways.

**Methods:** Low density TaqMan miRNA assays were used to quantify levels of 762 mature miRNAs under LIF stimulation in three choriocarcinoma-derived (JEG-3, ACH-3P and AC1-M59) and a trophoblast immortalized (HTR-8/SVneo) cell lines. Expression of selected miRNAs was confirmed in primary trophoblast cells and cell lines by qPCR. Targets and associated pathways of the differentially expressed miRNAs were inferred from the miRTarBase followed by a KEGG Pathway Enrichment Analysis. HTR-8/SVneo and JEG-3 cells were transfected with miR-21-mimics and expression of miR-21 targets was assessed by qPCR.

**Results:** A similar number of miRNAs changed in each tested cell line upon LIF stimulation, however, low coincidence of individual miRNA species was observed and occurred more often among choriocarcinoma-derived cells (complete data set at http://www.ncbi.nlm.nih.gov/geo/ under GEO accession number GSE130489). Altered miRNAs were categorized into pathways involved in human diseases, cellular processes and signal transduction. Six cascades were identified as significantly enriched, including JAK/STAT and TGFB-SMAD. Upregulation of miR-21-3p was validated in all cell lines and primary cells and STAT3 was confirmed as its target.

**Discussion:** Dissimilar miRNA responses may be involved in differences of LIF effects on trophoblastic cell lines.

## INTRODUCTION

Leukemia inhibitory factor (LIF) is a pleiotropic cytokine known to be indispensable for human reproduction. LIF controls uterine receptivity and influences trophoblast behavior by promoting proliferation, invasion and differentiation, and its dysregulation is associated with infertility and poor pregnancy outcome (reviewed in [1, 2]).

To investigate the molecular mechanisms involved in trophoblast response, several cell lines have been employed as models for primary cells [3–7]. Unlike primary cells, these models have the advantage to proliferate easily in culture while maintaining characteristics of desired trophoblast phenotypes. These models have been employed for decades, but it is still discussed how their limitations may influence translation of results to the *in vivo* situation and which markers are required to consider them as suitable trophoblast models [8]. Secreted proteins (e.g. IGF2, CSH1), trophoblast lineage markers (e.g. KRT7, VE-Cadherin, HLA-G) and expression of homeobox and imprinting genes (e.g. H19, IGF2, CDKN1C) are among the markers used to characterize trophoblast phenotypes [8, 9]. Recently, specific mRNA and microRNA (miRNA) expression profiles have been added to that list [10–12].

Cumulating evidence demonstrates that physiological characteristics of trophoblast biology (immunoregulation, invasion, proliferation among others) are tightly regulated by miRNAs [13–15]. These short non-coding RNA sequences (∼22 nt) regulate gene expression by sequence-specific binding to 3’UTR of mRNA targets resulting in their degradation or translational repression. We have previously reported differences in the miRNA expression profiles of four cell lines widely used as trophoblast models [10]: JEG-3, a human choriocarcinoma cell line preserving several trophoblast-like capacities; two hybrids of a JEG-3 mutant with human first and third trimester trophoblast cells, ACH-3P and AC1-M59 cells respectively [3–5]; and the immortalized human first-trimester trophoblast cell line HTR-8/SV40 [6]. Unlike primary trophoblast cells and choriocarcinoma cell lines, HTR-8/SV neo cells do not express the placenta-related Chromosome 19 MicroRNA Cluster (C19MC), but retain expression of C14MC [10]. In contrast, the mentioned choriocarcinoma-derived cell lines express C19MC, but only low levels of C14MC miRNA [10]. Likewise, these cell lines have differences in proliferation and invasion rates and respond dissimilarly under stimulation with LIF and other IL-6 family cytokines [16, 17]. Altogether, these data demonstrate that no cell line resembles all characteristics of primary trophoblast cells. Therefore, to understand the *in vivo* relevance of findings obtained by using these cell lines, comparison of several models is advised.

In order to gain more insights into the trophoblast response to LIF, four cell models were investigated to identify in common altered miRNAs, their associated pathways and targets. The LIF/STAT3/miR-21-3p axis was additionally validated in primary trophoblast cells.

## MATERIALS AND METHODS

*The Placenta Lab strictly applies quality management and is certified after DIN EN ISO 9001*.

### Primary trophoblast cells and cell lines

Healthy term placentae were obtained from the Department of Obstetrics, Jena University Hospital, Jena, Germany. Trophoblast isolation was performed using a modified Kliman method as described in detail elsewhere [10, 18]. Isolated cells have been checked for purity by flow cytometry using anti-EGF-receptor, anti-cytokeratin-7, and anti-HLA-G antibodies. The levels of impurity (cells negative for mentioned markers) did not exceed 10%.

JEG-3 cell line was purchased from DSMZ (Braunschweig, Germany). ACH-3P and AC1-M59 cells were a kind gift from Prof. G. Desoye (Graz, Austria). The immortalized cell line HTR-8/SV40 was kindly provided by Dr. Charles H. Graham, (Queen’ University, Kingston, Canada).

### Cell culture

Cell cultures were maintained under standard conditions (37°C, 5% CO_2_, humid atmosphere) in Ham’s F-12 Nutrient Mixture with L-glutamine (GIBCO) or RPMI Medium (GIBCO; HTR-8 cells) supplemented with 10 % heat-inactivated fetal calf serum (FCS; GIBCO) and 1 % penicillin/streptomycin antibiotic solution (GIBCO). Cells were checked regularly for mycoplasma contamination and their identity was confirmed by Short Tandem Repeat (STR) genotyping compared to the databanks of the American Type Culture Collection (ATCC) and the Leibniz Institute DSMZ – German Collection of Microorganisms and Cell Cultures (DMSZ).

### RNA isolation and miRNA TaqMan low density arrays

Cells were cultivated in 12-well plates and allowed to attach overnight. Subsequently, they were serum deprived for at least 2 h and then challenged 4 h with 10 ng/ml LIF (Millipore) [19, 20]. Total RNA was isolated using the mirVana isolation kit (Life Technologies), according to the manufacturer’s protocol. RNA purity was assessed by the ratio of spectrophotometric absorbance at 260 and 280 nm (A260/280nm) using NanoDrop ND-1000 (NanoDrop Inc). Total RNA (100 ng) was reverse transcribed using the specific Megaplex RT kit (Life Technologies) followed by a pre-amplification step. Expression of 754 miRNAs was profiled by using the TaqMan® Array Human MicroRNA A+B Cards Set v3.0 (Life Technologies) on a 7900HT Sequence detection system. Experimental data were analyzed using the DataAssist v3.0 Software (Life Technologies) using the formula 2^−ΔΔCt^ with RNU48 and RNU44 as endogenous controls.

### Validation of miRNA expression by qPCR

The expression levels of miR-885-5p, miR-21-3p and miR-34b-3p were verified by individual TaqMan miRNA Assays (Applied Biosystems, Foster City, CA, USA) according to the protocol provided by the supplier. Reverse transcription was performed with miRNA specific stem-loop RT primer (miR-21-3p Assay-ID 002438, miR-885-5p 002296, miR-34b 000427) using the TaqMan MicroRNA Reverse Transcription Kit (Applied Biosystems). qRT-PCR was done using the specific TaqMan Assay and TaqMan Universal PCR Master Mix no UNG. All reactions were run in duplicates including no-template controls in 96-well plates on a 7300 Real Time PCR System (Applied Biosystems). Fold changes were calculated using the formula 2^−ΔΔCt^ relative to non-stimulated cells with RNU48 as endogenous control.

### Prediction of targets and pathway analyses

Potential target genes of miRNA affected upon LIF stimulation (|FC| ≥ 1.5) were obtained from the miRTarBase [21] (release 7.0) to get non-redundant lists of target proteins for each of the different conditions. In the next step, all 299 available regulatory human pathways of the Kyoto encyclopedia of genes and genomes (KEGG) database [22] were ranked individually for the investigated conditions, according to the amount of targeted proteins within each pathway.

To evaluate whether a given pathway was overrepresented in the list of differentially affected miRNA targets, the hypergeometric test [23] was used to calculate corresponding p-values for each pathway:

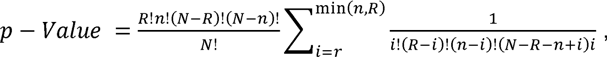

where *N* is the total number of miRNAs that have targets in any of the given pathways, *R* the number of miRNAs of interest (here miRNAs with an absolute fold change ≥ 1.5 upon LIF stimulation), *n* the total number of miRNAs with targets in the currently observed pathway and *r* the number of miRNAs of interest that have targets in the currently observed pathway.

After Benjamini-Hochberg false discovery rate (FDR) adjustment [24], only pathways with an enrichment p-value of ≤ 0.05 were considered statistically significant.

Gene Ontology analysis (GO) was used for identifying biological functions of the enriched KEGG pathways. Significantly altered pathways were organized within the seven networks of the KEGG pathway database [22]. Analysis of sub-pathways was performed by manual identification of altered key regions within the significantly enriched pathways.

### Validation of miR-21-3p targets

HTR-8/SVneo and JEG-3 cells were seeded in 6-well plates and allowed to attach overnight to reach 50% confluence at the time of transfection. Transfection was performed for 24 h using Oligofectamine (ThermoFisher) according to the manufacturer’s protocol as described before [25]. Sequences mimicking miR-21-3p (ID: MC12979) or a non-genomic scrambled sequence SCR (ID: MH12979) were transfected at a final concentration of 20 nM.

Expression level of STAT3 and SMAD7 was determined 24h after transfection by qPCR. RNA was isolated as described above. 100 ng were reverse transcribed using High-Capacity RNA-to-cDNA™ Kit (Applied Biosystems, Darmstadt, Germany). Quantitative real-time PCR was performed using TaqMan assays (STAT3, Assay ID: Hs00374280_m1; SMAD7, Assay ID: Hs00998193_m1 and GAPDH, Assay ID: Hs02758991_g1) and TaqMan Universal PCR Master Mix reagents (Applied Biosystems). qPCR was run on a Mx3005P qPCR System (Applied Biosystems). Expression of STAT3 and SMAD7 was normalized using the 2^−ΔCt^ or 2^−ΔΔCt^ method relative to GAPDH and the respective experimental control.

## RESULTS

### miRNA profile after LIF stimulation changes in a cell-specific manner

After 4 h of LIF treatment (10 ng/ml), 754 miRNAs were profiled and miRNAs with Ct ≤ 35 displaying more than 1.5-fold change were considered altered. In HTR-8/SVneo cells, expression of 110 miRNAs was influenced (39 up- and 71 down-regulated). In the choriocarcinoma-derived JEG-3 cell line, 91 affected miRNAs were identified: 45 down- and 36 up-regulated. In the hybrid cell lines AC1-M59 and ACH-3P cells, a total of 88 (45 down- and 43 up-regulated) and 130 (45 down- and 85 up-regulated) were identified, respectively (Figure 1). The microarray dataset is available at the Gene Expression Omnibus database http://www.ncbi.nlm.nih.gov/geo/ under GEO accession number GSE130489.

**Figure 1.**
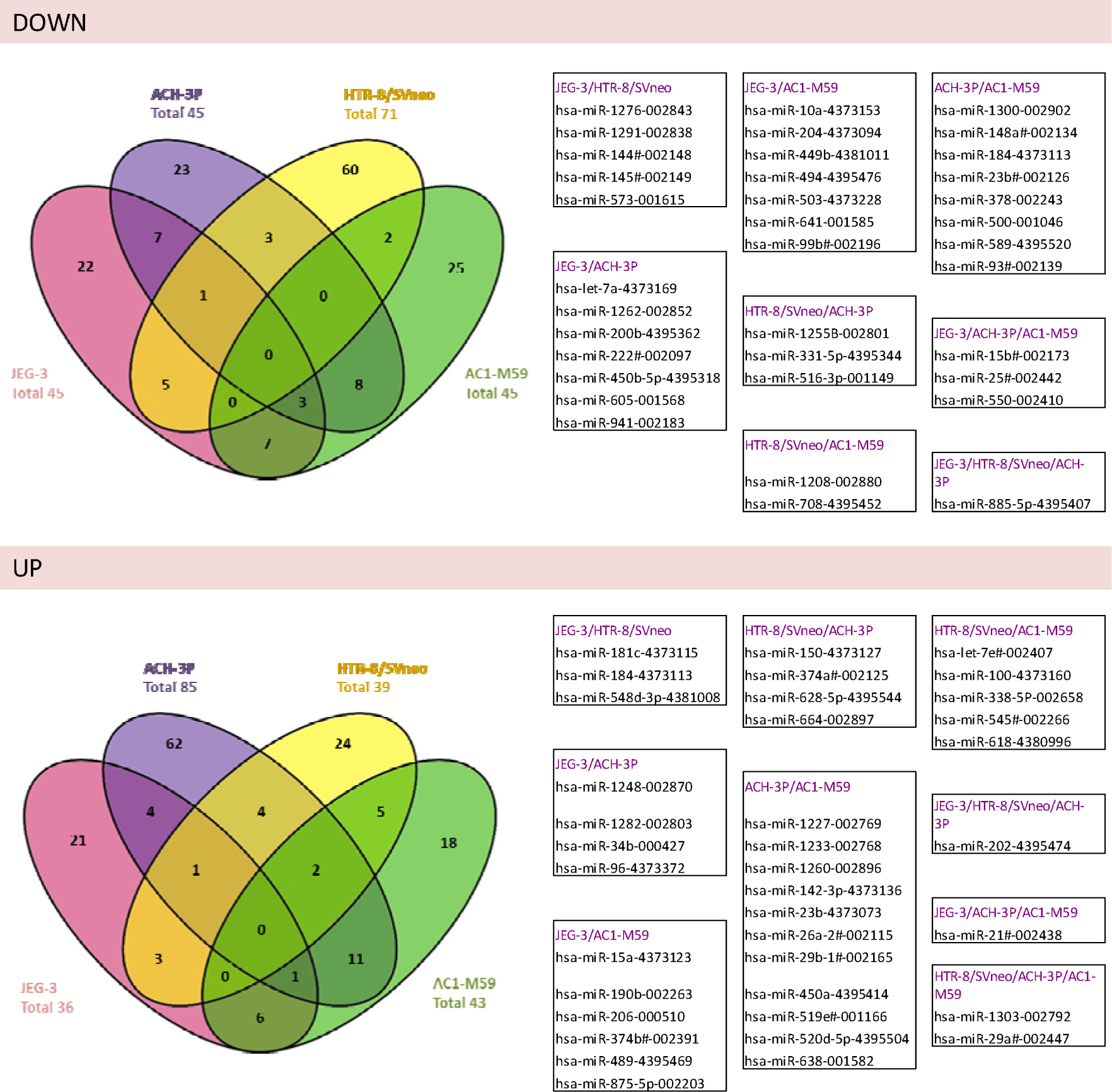
miRNAs in trophoblastic cells regulated upon LIF treatment. Venn diagrams show altered miRNAs (mean fold-change ≥ 1.5; n=3) in cells treated with LIF (10 ng/ml; 4h) compared to controls. miRNAs in boxes were altered in at least two cell lines. miRNA species are reported as hsa-miR-Number-AssayID.

Under the selected conditions for the data analysis of the arrays, no miRNA species was identified simultaneously altered in all four tested cell lines. ACH-3P and AC1-M59 cells had the most similar response with 25 altered miRNAs in common (14 up- and 11-down regulated), followed by pairs of JEG-3:ACH-3P and JEG-3:AC1-M59 cells, each with 17 shared altered miRNAs (Figure 1).

To confirm the results obtained from the arrays and to validate the observed effects on primary cells, three miRNAs were selected for further analysis. Both miR-885-5p and miR-21-3p were identified altered in three cell lines and their basal expression in primary trophoblast cells resembles that of HTR-8/SVneo and choriocarcinoma cell lines [10]. Additionally, miR-34b-3p was included for verification by qPCR as it was found altered only in two cell lines. Although having a tendency to decrease upon LIF stimulation in all tested models, reduction of miR-885-5p was only significant in JEG-3 and ACH-3P cells. miR-21-3p was significantly up-regulated in all tested cell lines and in primary trophoblast cells. No significant changes were detected in miR-34b-3p (Figure 2).

**Figure 2.**
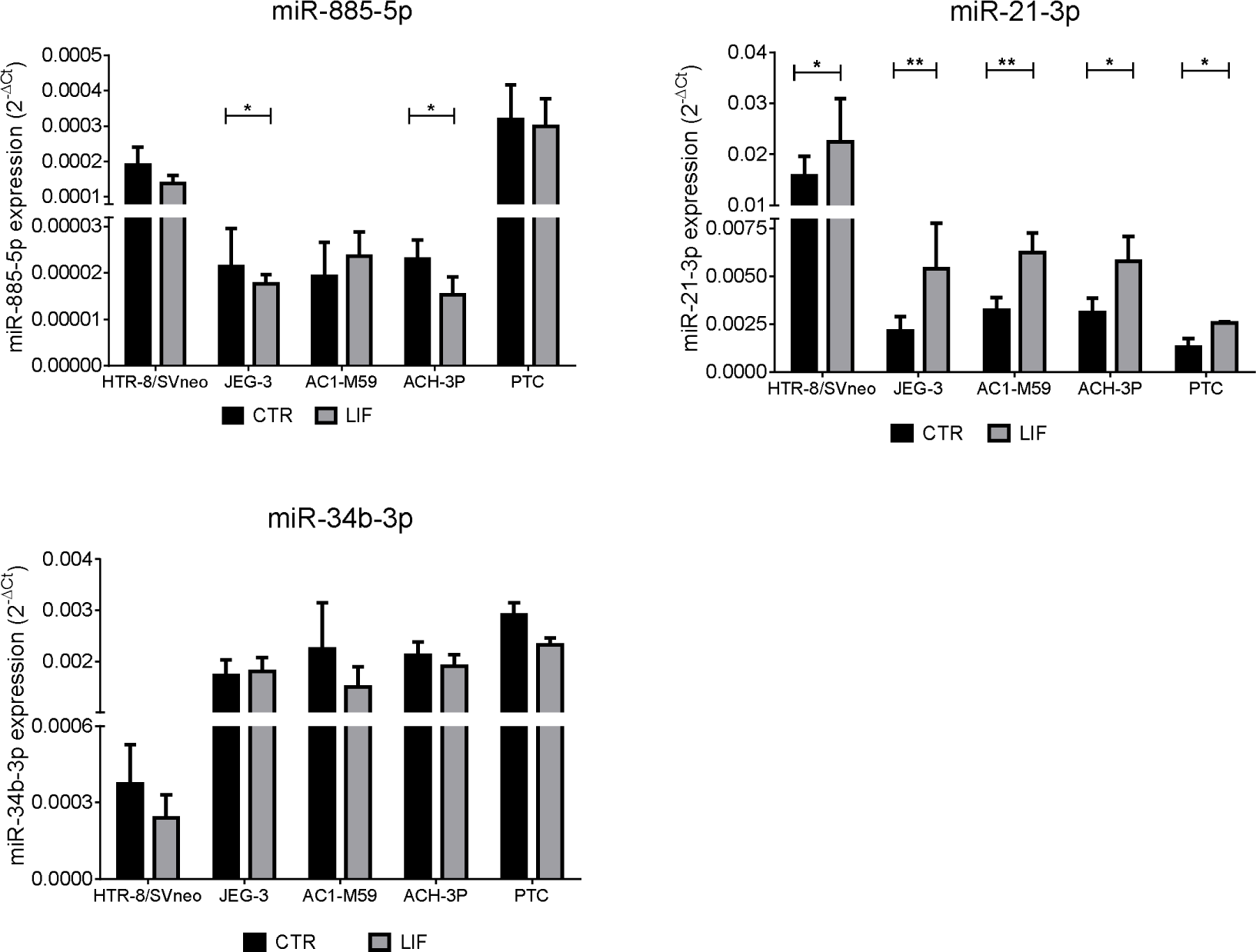
Verification of miRNA array expression data in primary cells. Expression of miRNAs identified as altered in several cell lines upon LIF treatment was confirmed by qPCR in cell lines and primary trophoblast cells (PTC). Fold change expressed as 2^−ΔCT^ to RNU48. Values are presented as mean ±SEM (n = 4), * p < 0.05, ** p< 0.01 (vs.CTR)

### Gene oncology and KEGG pathway analysis of the affected miRNAs

Targets of the differentially expressed miRNAs were inferred from the miRTarBase release 7.0. All 299 available regulatory human pathways of the KEGG database were ranked individually according to the amount of targeted proteins after LIF stimulation within each pathway.

Only two pathways were identified in JEG-3 as most likely to be affected after LIF treatment: Epithelial cell signaling in Helicobacter pylori infection hsa05120 and Adherens junction hsa04520. In ACH-3P, AC1-M59 and HTR-8/SVneo cells, 17, 41 and 31 statistically significantly enriched pathways were identified, respectively (Suppl. Figure 1). Among them, a group of 13 KEGG pathways were found significantly enriched in at least two cell lines (Suppl. Figure 1).

### Analysis of subpathway regions and associated functions

To further establish conserved pathways activated by LIF in trophoblastic cell lines, local areas of high concentration of target genes were identified within high ranked pathways and designated as sub-pathways. Six sub-pathways were identified as influenced by LIF affected miRNAs: a) NFkB, b) TGFB-Smad, c) RAS-RAF-MEK-ERK, d) PI3K-Akt, e) NWASP-ARP2/3-(F-)actin and f) JAK/STAT. Among them, NWASP-ARP2/3-(F-)actin was identified altered in all tested cell lines. Alterations of a, c, d and f were identified in ACH-3P, HTR-8/SVneo and AC1-M59 cells, and of TGFB-Smad in JEG-3 and HTR-8/SVneo cells (Figure 3A and 3B). Additionally, a cross-talk between components of subpathways (a, c, d and f) was also described (Figure 3A).

**Figure 3.**
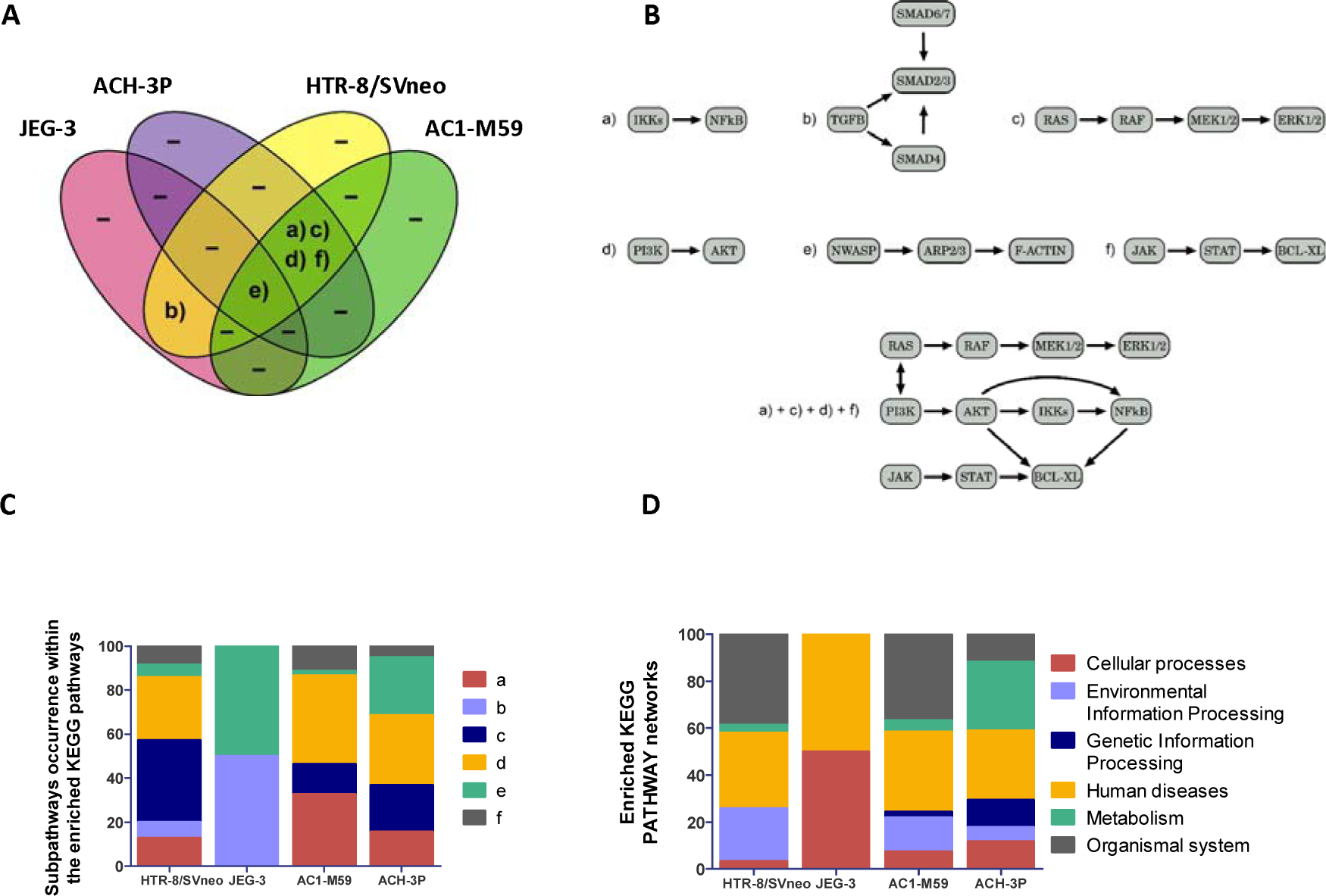
Subpathways and biological functions of miRNAs modulated by LIF. A) Venn diagram depicts the identified modulated subpathways and how they are shared between cell lines. B) Identified subpathways a – NFkB activation by members of the IKK complex; b – TGF-beta/SMAD pathway; c – Ras-Raf-MEK-ERK pathway; d – PI3K/AKT/mTOR pathway; e – N-WASP and Arp2/3 regulation of actin; f – JAK/STAT signaling pathway. Known interactions between four of those subpathways are also depicted. C) Overview of the relative number of occurrences of the identified subpathways within the significantly enriched Kyoto Encyclopedia of Genes and Genomes (KEGG) pathways. D) Putative biological processes affected by LIF based on the KEGG orthology reference hierarchy.

Occurrence analysis revealed that some subpathways were more frequently altered across enriched pathways resulting in different profiles among cell lines (Figure 3C). Enrichment of PI3K/AKT was observed in HTR-8/SVneo, ACH-3P and AC1-M59 cells, while RAS/RAF enrichment was also frequent in HTR-8/SVneo cells (Figure 3C).

Finally, to identify biological functions of LIF in trophoblast cells, significantly altered pathways were organized within the KEGG categories. Those which were identified in all cell lines were “human diseases” and “cellular processes” (Figure 3D).

### Validation of putative targets of miR-21-3p

miRTarBase (release 7.0) [21] predicts STAT3 and SMAD7 expression as affected by the LIF-induced increase of miR-21-3p (Figure 4A). After challenging with LIF, qPCR showed that STAT3 mRNA was significantly increased in all tested cell lines and in primary trophoblast cells (Figure 4B). SMAD7 was altered in JEG-3 but not in HTR-8/SVneo cells and was therefore not considered for further analysis (data not shown). Overexpression of miR-21-3p in HTR-8/SVneo and JEG-3 cells resulted in a significant decrease of STAT3 mRNA confirming it as target (Figure 4C).

**Figure 4.**
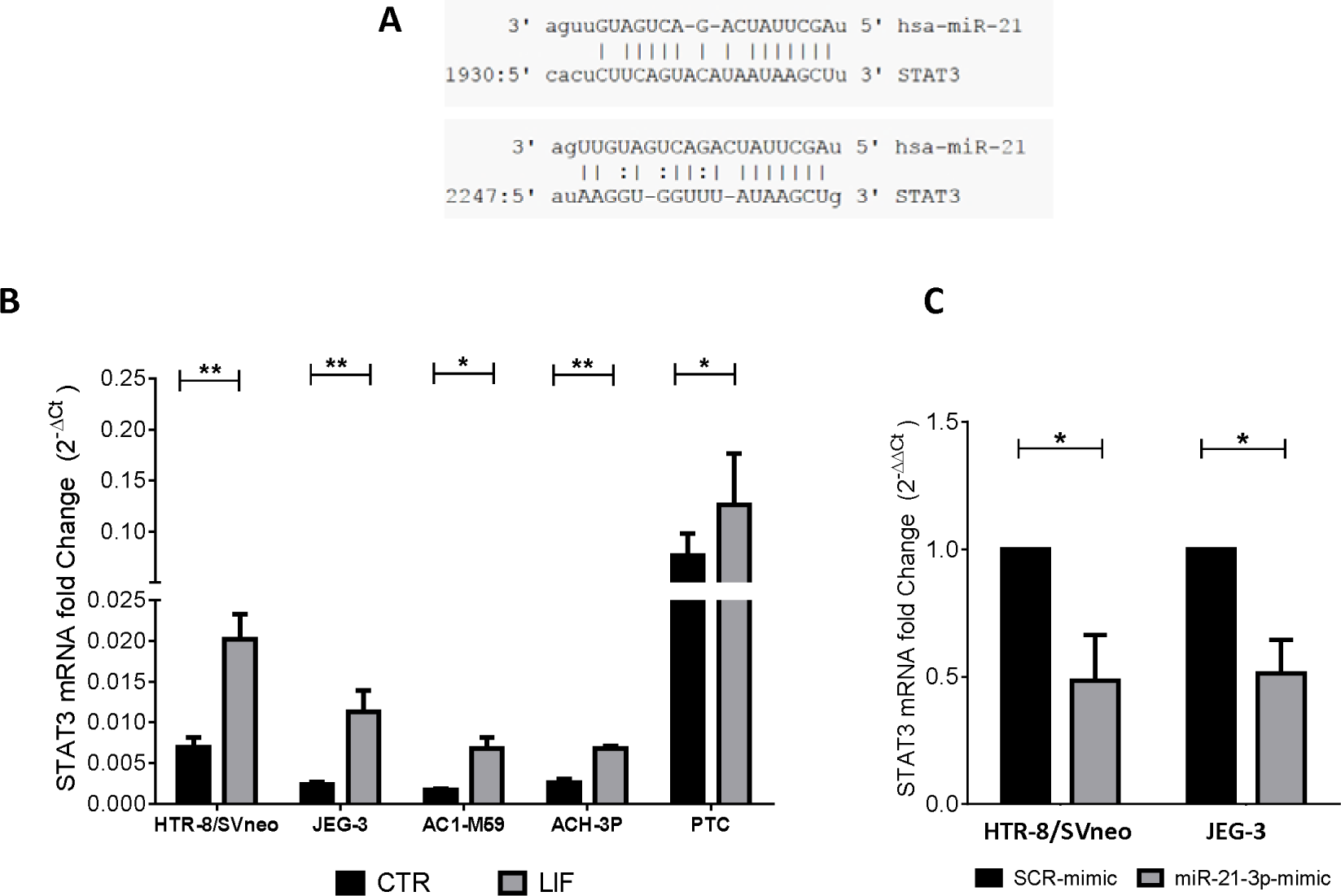
miR-21-3p targets STAT3 in trophoblast cells. A) miRNA/mRNA alignment of miR-21-3p/STAT3. B) Expression of STAT3 mRNA upon LIF stimulation (10 ng/ml; 4h) in cell lines and primary trophoblast cells (PTC). C) Expression of STAT3 mRNA in HTR-8/SVneo and JEG-3 cells after transfection with SCR- and miR-21-3p-mimics. Data expressed as 2^−ΔCT^ or 2^−ΔΔCT^ to GAPDH. One-way ANOVA test *p < 0.05 **p<0.01.

## DISCUSSION

Leukemia Inhibitory Factor (LIF) is one of the most important factors within the uterine environment. It is involved in fundamental processes for human reproduction including endometrium receptivity and blastocyst implantation [2]. LIF and its receptor LIFR are localized in the chorionic villi, decidua and extravillous trophoblast cells (EVTs) of first trimester and term placenta [26]. There, LIF exerts an important role in the regulation of EVT invasion and the communication of trophoblast with decidua and immune cells, which is required for an adequate spiral artery remodeling [27]. Activation of several signaling pathways by LIF and the resulting biological functions in trophoblastic cells have been described previously by our group and others [17, 28–33]. However, miRNAs involved in these processes and the similarities across different trophoblastic models are not known.

To circumvent the disadvantages of working with primary cells and still allowing the study of intracellular signals, several cell lines have been established to investigate functions on trophoblastic cells. Some of the most widely used models have been derived from choriocarcinoma (e.g. JEG-3, ACH-3P and AC1-M59) or from primary cells immortalized with the gene encoding for simian virus 40 large T antigen (HTR-8/SVneo) [3–6]. Due to the differences in their origin, these models exhibit unique expression patterns of genes, proteins [12, 34] and miRNAs [10]. These differences are reflected in their ability to proliferate, migrate and invade, thus, constituting an important aspect when selected as suitable models for trophoblast research.

Rather than to authenticate the “most suitable” trophoblast cell line for human research, this study aimed to determine the most affected miRNAs and their associated pathways involved in the response of trophoblast cells to LIF. Expression of 754 miRNAs was investigated by miRNA arrays revealing a similar number of altered miRNAs in all four analyzed cell lines upon stimulation with LIF. Remarkably, overlapping of specific miRNAs was low and occurred mostly between choriocarcinoma cell lines. This may rely on the shared origin of JEG-3, AC1-M59 and ACH-3P cells, but it also may be the result of miRNA redundancy in HTR-8/SVneo cells to compensate lack of some placenta-related miRNAs (e.g. C19MC miRNAs) [10]. However, further investigation should confirm this assumption.

Identification of affected miRNAs was performed under a strict threshold of |fold change| ≥ 1.5 and an expression of Ct ≤ 35. We consider that these strict selection criteria may also contribute to the low coincidence of results among cell lines, but this strategy allows the identification of the most relevant changes while avoiding false positive results. Likewise, a single time point (4h) and a unique concentration (10 ng/ml) were selected for this study based on our previous reports on LIF functions and LIF-mediated alterations of miRNAs, mRNAs and proteins in trophoblastic cells [17, 28–30]. Due to this selection, differences in the kinetic and sensitivity of the cell lines to LIF may be disregarded resulting in oversight of additional shared miRNAs.

Further, the miRNA arrays used in this study allow the simultaneous measurement of hundreds of miRNA species, but this technique may neglect the variation in melting temperatures of miRNA probes and the results depend greatly on the RNA input and normalization [35]. Thus, qPCR is still considered the “gold standard” for the corroboration of results obtained from miRNA arrays. We selected three miRNAs for verification based on their similarities in the basal expression to primary trophoblast cells. E.g. in primary cells expression of miR-885-5p was similar to that of HTR-8/SVneo cells whilst expression of miR-21-3p and miR-34b-3p was closer to that of choriocarcinoma-derived cell lines [10]. Approximately, 67% of the results in the arrays were confirmed by qPCR which is consistent with other studies [35]. Furthermore, miR-21-3p, which was identified upregulated in three cell lines by using the miRNA arrays, was confirmed upregulated in all cell lines and in primary trophoblast cells by qPCR. Previously, we reported upregulation of the miR-21 guide strand (miR-21-5p) in trophoblastic cells stimulated with LIF [28]. miR-21-5p correlates positively with the ability of trophoblastic cells to invade and proliferate, but its targets and associated pathways differ between JEG-3 and HTR-8/SVneo cells [13]. The function of miR-21-3p on trophoblast cells remains to be determined but it is to hypothesize that it also plays a role in controlling cell proliferation and invasion.

To further examine the biological relevance of the array results, an *in silico* analysis was carried out. This strategy has been employed in other studies to assign functional meaning to regulation at mRNA level in trophoblast cells [11, 36]. In our study, upon LIF stimulation, targets of the differentially expressed miRNAs were firstly predicted by miRTarBase [21] and then mapped to KEGG pathways. To avoid a biased selection of the most relevant pathways associated to LIF-regulated miRNAs, all annotated KEGG pathways were included in the study without filtrating for those directly related to trophoblast cells. These pathways were then ranked based on the hit numbers and using the hypergeometric test to identify significantly altered pathways. Once more, the resulting pathways diverge among cell lines. Despite the similarities observed in single miRNA species, analysis of ACH-3P and AC1-M59 cell lines resulted in dissimilar enriched pathways. Likewise, while in JEG-3 only two pathways were significantly altered upon LIF stimulation, in AC1-M59 41 affected pathways and HTR-8/SVneo cells 31 pathways were identified.

Some of the enriched pathways belong to the KEGG DISEASE and KEGG DRUG databases [22] and thus, they may not necessarily have been described in the context of trophoblast behavior or reproduction. For instance, “Estrogen signaling pathway has04915” and “JAK-STAT signaling hsa04630” pathways significantly enriched in HTR-8/SVneo cells have been respectively identified in endometrium during the implantation period [37] and already recognized as a pivotal cascade for signaling of IL-6 cytokines in trophoblast cells [38]. Nevertheless, other apparently unrelated pathways were also identified such us “Alcoholism hsa05034” and “Non-alcoholic fatty liver disease (NAFLD) hsa04932” with low p-values in HTR-8/SVneo and AC1-M59, respectively.

To improve the interpretation of biological phenomena associated to enriched KEGG pathways, analysis of local regions or sub-pathways has been recently proposed [39]. By applying this method, mapped regions of high enrichment can be manually identified within the pathways of the previous step highlighting the core components targeted by specific miRNAs. For instance, LIF-altered miRNAs target in the pathway “Alcoholism” the “RAS-RAF-MEK-ERK1/2” subpathway, whilst in “NAFLD” the “IKK-NFkB” and “PI3K-Akt” subpathways. Using the same method for all enriched pathways, six sub-pathways were consistently mapped across cell lines: NFkB, TGFB-SMAD, RAS-RAF-MEK-ERK1/2, PI3K-Akt, NWASP-ARP2/3-(f)-actin and JAK-STAT. Because most of the enriched pathways can be organized within these sub-pathways, it can be suggested that they may be major cascades involved in the LIF response of trophoblast cells. Indeed, activation of JAK/STAT, MAPK and PI3K/Akt in trophoblast cells upon LIF stimulation has been described previously [17, 29, 30, 33].

Most of the enriched pathways identified by the *in silico* analysis were categorized into the KEGG PATHWAY networks of “Cellular processes” and “Human diseases” [22]. LIF is known as one of the most important factors mediating trophoblast biology [1, 2, 17, 32] and its dysregulation has been reported in pregnancy disorders such as preeclampsia [33] and hydatidiform mole [40]. Altogether, our results agree with those reports and contribute to identify the miRNAs and their associated pathways involved in those processes.

Finally, STAT3, an inductor of cell growth and invasion [41], and also pivotal regulator of trophoblast behavior [17, 41] was identified as putative target of miR-21-3p. We demonstrate increased expression of STAT3 mediated by LIF in all cell lines and in primary cells and confirmed it as target of miR-21-3p using two different cell lines. A recent report has shown based on bioinformatic analysis followed by ChIP-PCR that STAT3 acts as a transcription factor initiating miR-21 expression [42]. Our results indicate that by reducing STAT3 expression miR-21-3p may be part of the complex negative feedback system of STAT3.

Although the specific function of the miR-21-3p strand remains to be determined, the common response in the models suggests the LIF/STAT3/miR-21-3p axis to contribute in controlling major trophoblast functions.

In previous reports, we demonstrated that cellular behavior upon LIF treatment differs among trophoblastic cell lines. For instance, invasion is induced by LIF in JEG-3 and ACH-3P but not in AC1-M59 or HTR-8/SVneo cells [17, 29]. Now, we have demonstrated that these dissimilar LIF responses of trophoblastic cells may be influenced by the cell-specific set of altered miRNAs. However, these alterations converged in six major conserved pathways demonstrating the redundancy of miRNA species and highlighting the importance of further studying these pathways. Because the investigated cell lines have particular features and responses, they may reflect aspects of discrete populations of trophoblast cells present *in vivo*. Thus, their applicability as trophoblast models should be considered depending on the question to be addressed.

Altogether, this study demonstrates that LIF induces responses in miRNA expression in trophoblastic cells and reveals further differences in the behavior of the most frequently used trophoblast model cell lines.

## ACKNOWLEDGEMENT

The project has been supported by the German Research Foundation (DFG, Ma1550/12-1 to URM and RRF; Mo2017/3-2 to DMMP; MA 5082/7-1 to MM). It has been partially financed by the ProExzellenz program of the Thuringian Ministry for Research (EB: TMWWDG; RegenerAging – FSU-I-03/14).

---

**Suppl. Figure 1.**
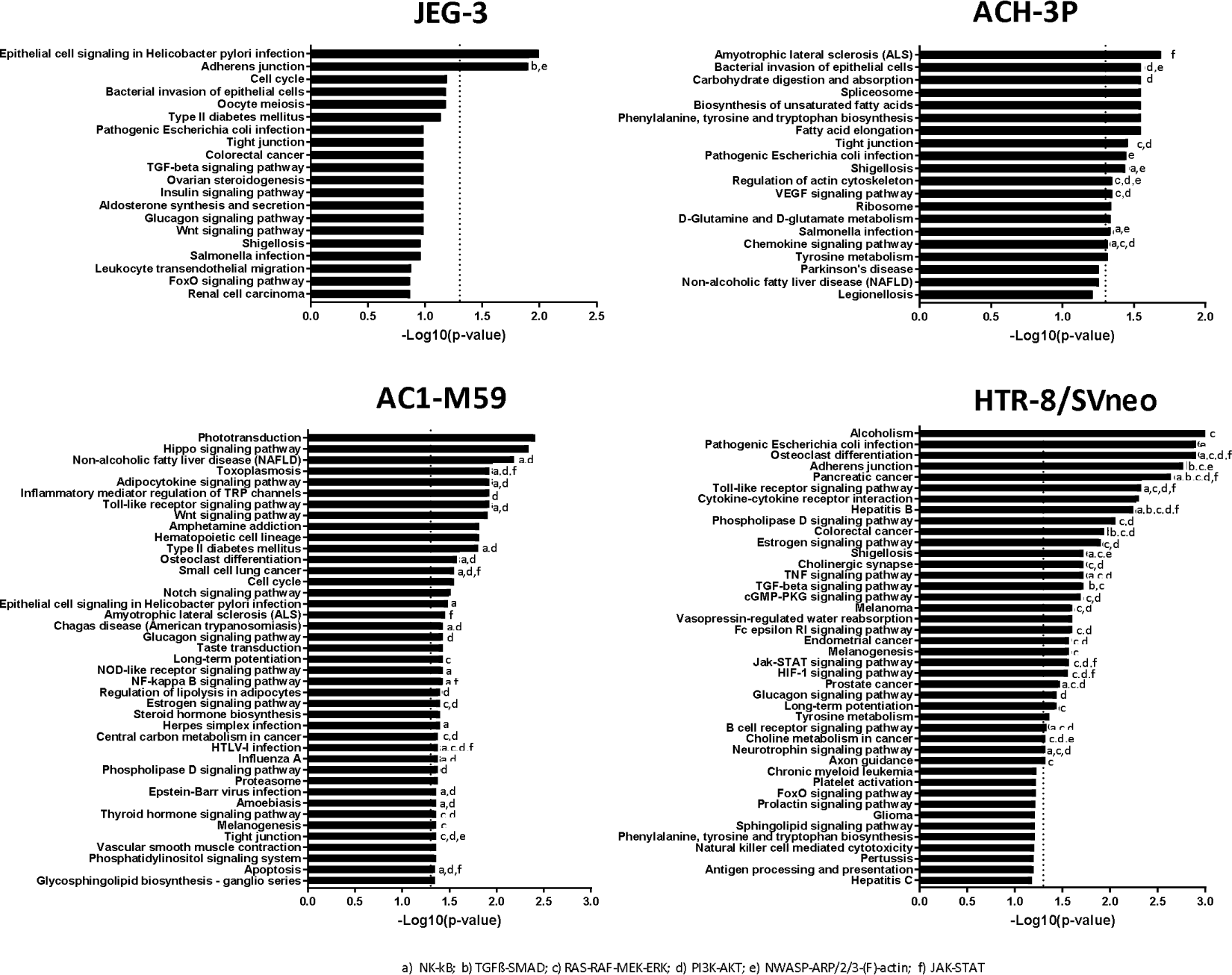
Signal pathway analysis of the target genes of LIF-altered miRNAs. Gene targets of miRNAs changing after LIF were inferred by TargetScan and classified into Kyoto Encyclopedia of Genes and Genomes (KEGG) pathways. Pathways were considered as significantly altered when p<0.05. Letters represent key subpathway regions a) NK-kB, b) TGFß-Smad, c) Ras-Raf-MEK-ERK, d) PI3K-Akt, e) NWASP-ARP/2/3-(F)-actin, f) JAK-STAT

